# Integrated metabolome and transcriptome analyses provide insight into colon cancer development by the gut microbiota

**DOI:** 10.1101/2020.07.24.220269

**Authors:** Susheel Bhanu Busi, Zhentian Lei, Lloyd W. Sumner, James Amos-Landgraf

**Affiliations:** University of Missouri School of Medicine, One Hospital Drive, MA204, Columbia, MO 65212; University of Missouri School of Veterinary Medicine, 1600 Rollins Street, Columbia, MO 65211; University of Missouri Division of Biochemistry, 117 Schweitzer Hall, Columbia, MO 65211; University of Missouri Metabolomics Center, 243 Christopher S. Bond Life Sciences Center, 1201 Rollins Street, Columbia MO 65211; Rat Resource and Research Center, University of Missouri, 4011 Discovery Drive, Columbia MO 65201

**Author notes:** **Corresponding Author:** James Amos-Landgraf, University of Missouri School of Veterinary Medicine, 1600 Rollins Street, Columbia, MO 65211.

**Keywords:** colon cancer, metabolomics, transcriptomics, gut microbiota and integrated analyses

## Abstract

Colon cancer onset and progression is strongly associated with the presence, absence, or differences in relative abundances of certain microbial taxa in the gastrointestinal tract. However, specific mechanisms affecting disease susceptibility related to complex commensal bacterial mixtures are poorly understood. We used a multi-omics approach to determine how differences in the complex gut microbiome (GM) influence the metabolome and host transcriptome and ultimately affect susceptibility to adenoma development in a preclinical rat model of colon cancer. Fecal samples from rats harboring distinct complex GMs were analyzed using ultra-high performance liquid chromatography mass spectrometry (UHPLC-MS). We collected samples prior to observable disease onset and identified putative metabolite profiles that predicted future disease severity, independent of GM status. Transcriptome analyses performed after disease onset from normal epithelium and tumor tissues between the high and low tumor GMs suggests that the GM is correlated with altered host gene expression. Integrated pathway (IP) analyses of the metabolome and transcriptome based on putatively identified metabolic features indicate that bile acid biosynthesis was enriched in rats with high tumors (GM:F344) along with increased fatty acid metabolism and mucin biosynthesis. These data emphasize the utility of using untargeted metabolomics to reveal signatures of susceptibility and resistance and integrated analyses to reveal common pathways that are likely to be universal targets for intervention.

**Statement of significance:** Fecal metabolites, influenced by the gut microbiota, correlate with colon adenoma risk in a preclinical model of familial colon cancer.

## Introduction

Colorectal cancer is the second leading cause of cancer death and remains difficult to diagnose without invasive or non-universally available procedures such as colonoscopy (1). Several recent studies in animal models and human patient populations have begun to identify biomarkers that have some diagnostic capability (2–4). Additionally, association studies in patient populations have shown positive and negative correlations with various bacterial species (5, 6). The association between certain bacterial species and a quantifiable impact on colon adenoma development has also been shown in animal models (7, 8). However, the link between early diagnostic biomarkers and the gut microbiota has not been sufficiently investigated and the mechanisms driving phenotypic differences are not well determined. The diagnostic ability for biomarkers to correlate with early colon cancer is likely owing, at least in part, to bacterially-derived metabolites and the corresponding host responses to these metabolites (9–11).

Untargeted metabolomics is a maturing field focused on the large-scale quantitative and qualitative analyses of small molecular weight (<2000) biomolecules. Information from these studies provide unique insight into physiological pathways that have important roles in health and disease (12). Given that microbial species play a critical role in both production and use of host metabolites, it is likely that the GM has a substantial impact on the overall metabolite composition (13, 14). Confirming this hypothesis, studies have demonstrated significant differences in metabolites between germ-free mice and their conventionally housed counterparts, emphasizing a microbiota-driven metabolic profile (15). As a result, the role of metabolic mediators as intermediates between the GM and tumorigenesis in both rodent models and humans has garnered substantial interest. Dazard *et al*. used mass spectrometry to determine that plasma from *Apc^Min^* mice had a distinct metabolome compared to wild-type (WT) littermates (16). However, due to a lack of longitudinal metabolomics data in this study and others, it is unclear whether these metabolic changes are a consequence of tumor development or are causative of tumor initiation or progression.

We previously reported that naturally occurring gut microbiota (GM) can modulate colon cancer susceptibility in a preclinical rat model of Familial Adenomatous Polyposis. We rederived isogenic embryos of the F344/NTac-Apc^+/*Pirc*^ rat model into different populations of surrogate dams each harboring distinct gut microbiota: GM:F344, GM:SD and GM:LEW. Through this method we created animals that harbored distinct endogenous complex GMs. Pirc rats with the GM:F344 had the highest tumor burden, while GM:LEW rats had a significantly reduced tumor burden, including two animals that had no visible colonic adenomas at 6 months of age (7). The GM and metabolome separately have been shown to affect colon cancer tumorigenesis, however, there is insufficient data demonstrating how host gene expression is affected by the interplay of the microbiome and intestinal metabolites. We used a multi-omics approach to evaluate how differences in the microbiome affect the fecal metabolome and host gene expression to identify putative indicators of risk for adenoma development and begin to reveal the mechanisms by which the GM modulates disease susceptibility.

## Results

### Metabolite features at 1-month of age predict tumor susceptibility and severity at later developmental stages

Fecal samples collected from rederived Pirc rats harboring distinct GMs were analyzed by UHPLC-MS and found an average of 499 raw features of which 246 were shared among all samples and were used for subsequent analyses (Supplementary Table 1). Principal component ordination analysis (PCA) indicated a separation of 33.2% along the first component (PC1) accounting for some of the variability between each group (Fig.1A). Non-metric dimensional scaling also showed a similar separation (Fig. 1B) between groups, suggesting that the features identified via UHPLC-MS differentiated the high- (GM:F344) and low-susceptibility (GM:LEW) gut microbiota profiles, with GM:SD occupying the intermediate ordinates. Hierarchical clustering was performed using Euclidean distance and Ward’s clustering algorithm on the metabolomics dataset to identify the dissimilarity of the samples and groups with respect to each other. The Dendrogram demonstrates the separation of the metabolic profile of the fecal samples at 1 month, based on colonic tumor burden assessed at 6 months of age (Fig.1C).

**Fig.1.**
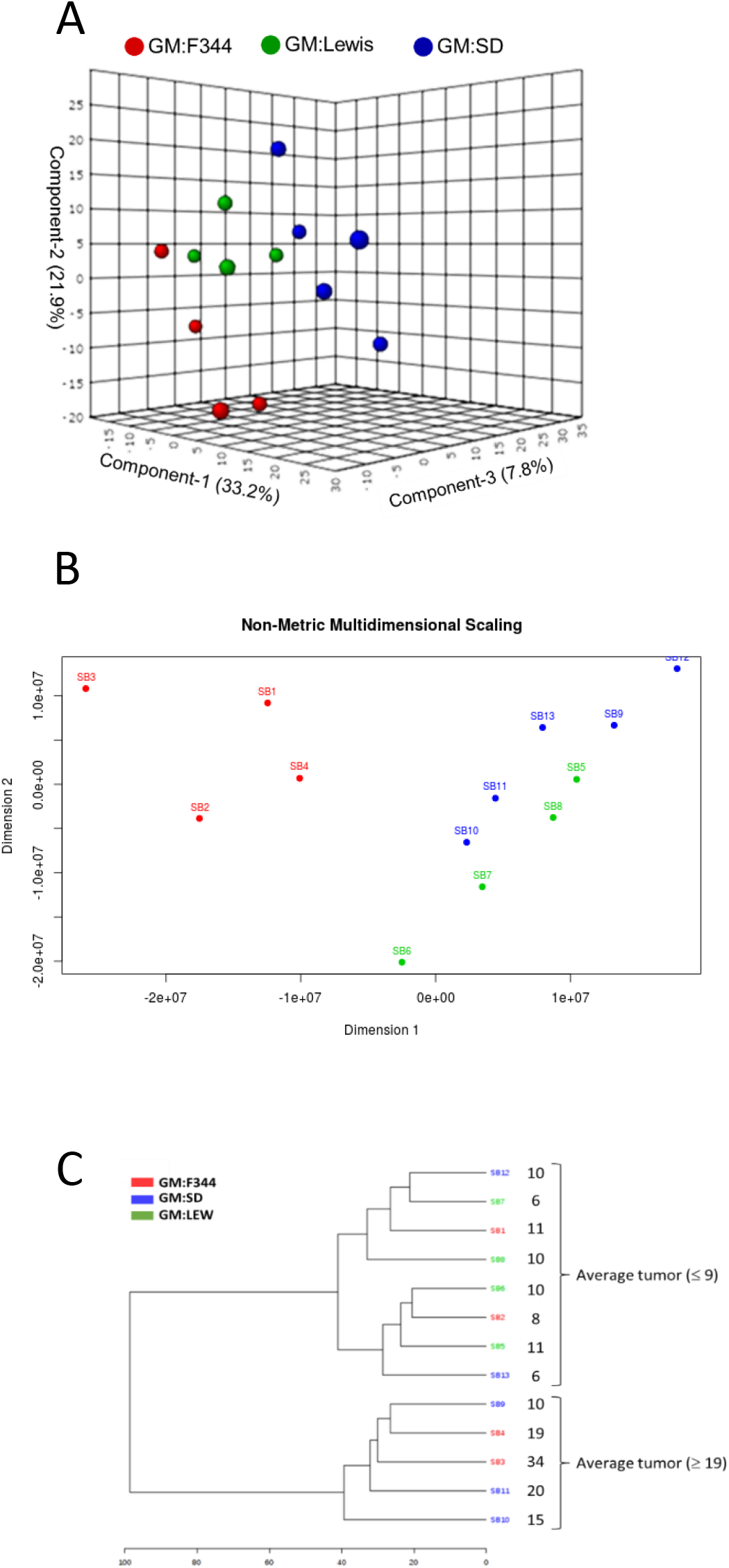
Metabolomics analyses indicate differential features in feces from differing gut microbiota

### Metabolomics analyses indicate differential metabolic profiles between GM:F344 and GM:LEW

Observing that GM:F344 and GM:LEW had the highest and lowest average number of tumors respectively, we further analyzed the differential features contributing to disease susceptibility within these groups using clustering and individual correlation analyses (Fig.2A and 2B). Using linear discriminant analyses (LDA) we identified the putative metabolites contributing to the high (GM:F344) and low (GM:LEW) tumor groups’ separation observed in the Dendrogram (Fig.2C). Some of the putative metabolites identified in the low tumor group, i.e. GM:LEW, showed up to a 4-fold increase compared to GM:F344 high tumor group (Fig. D). Tandem MS analysis was used to further identify the compounds that were differential between the low and high tumor groups. We generated tandem MS spectra for the compounds with the mz/rt values of 329.10/9.2 min and 315.12/6.39 min; however, their identities could not be definitively established based on the spectral libraries currently available. Comparison of the features without regard to specific GM and terminal tumor counts found significant correlations between individual metabolites at 1-month of age and terminal colonic tumor numbers (Fig.2E).

**Figure 2.**
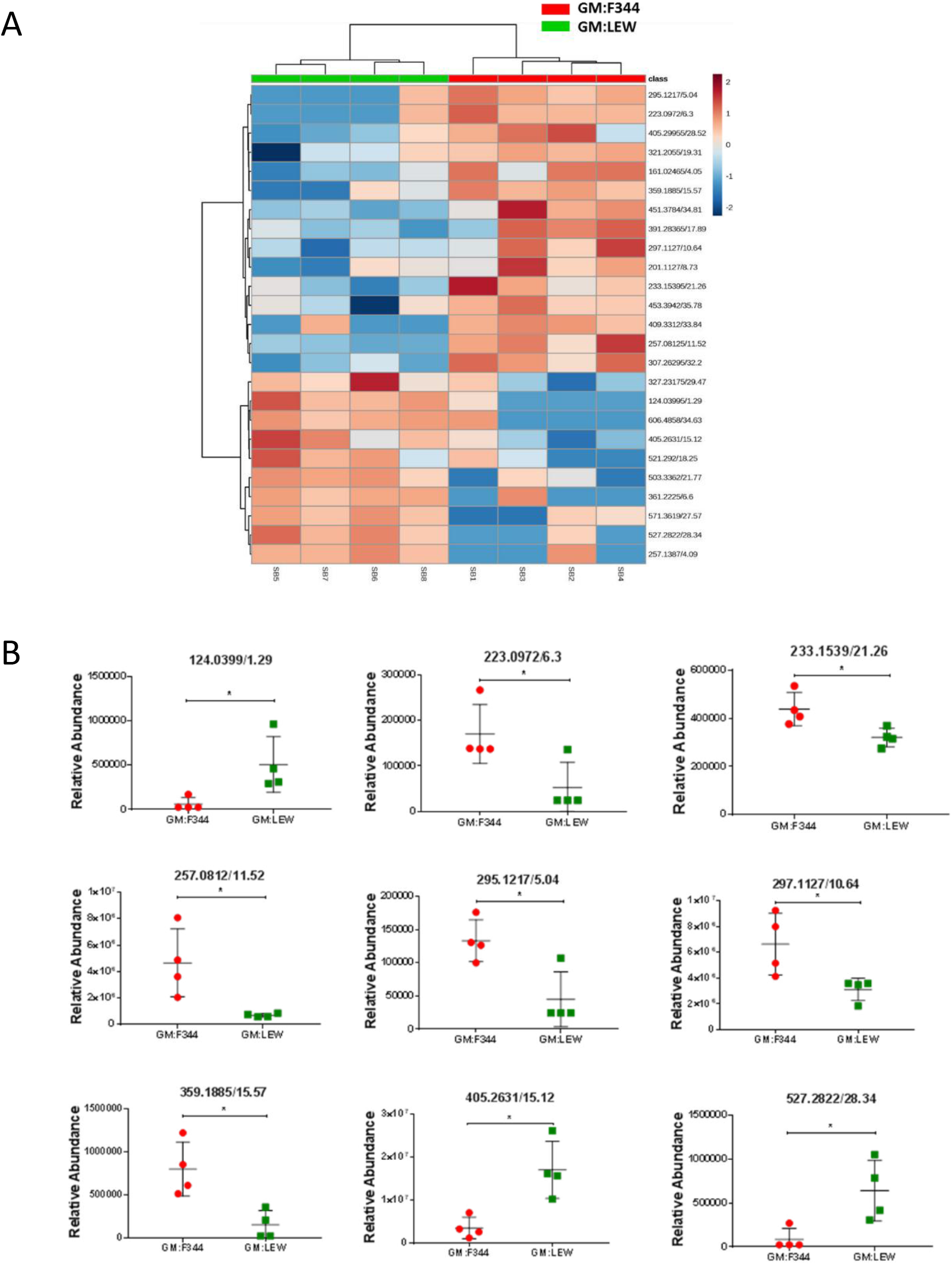
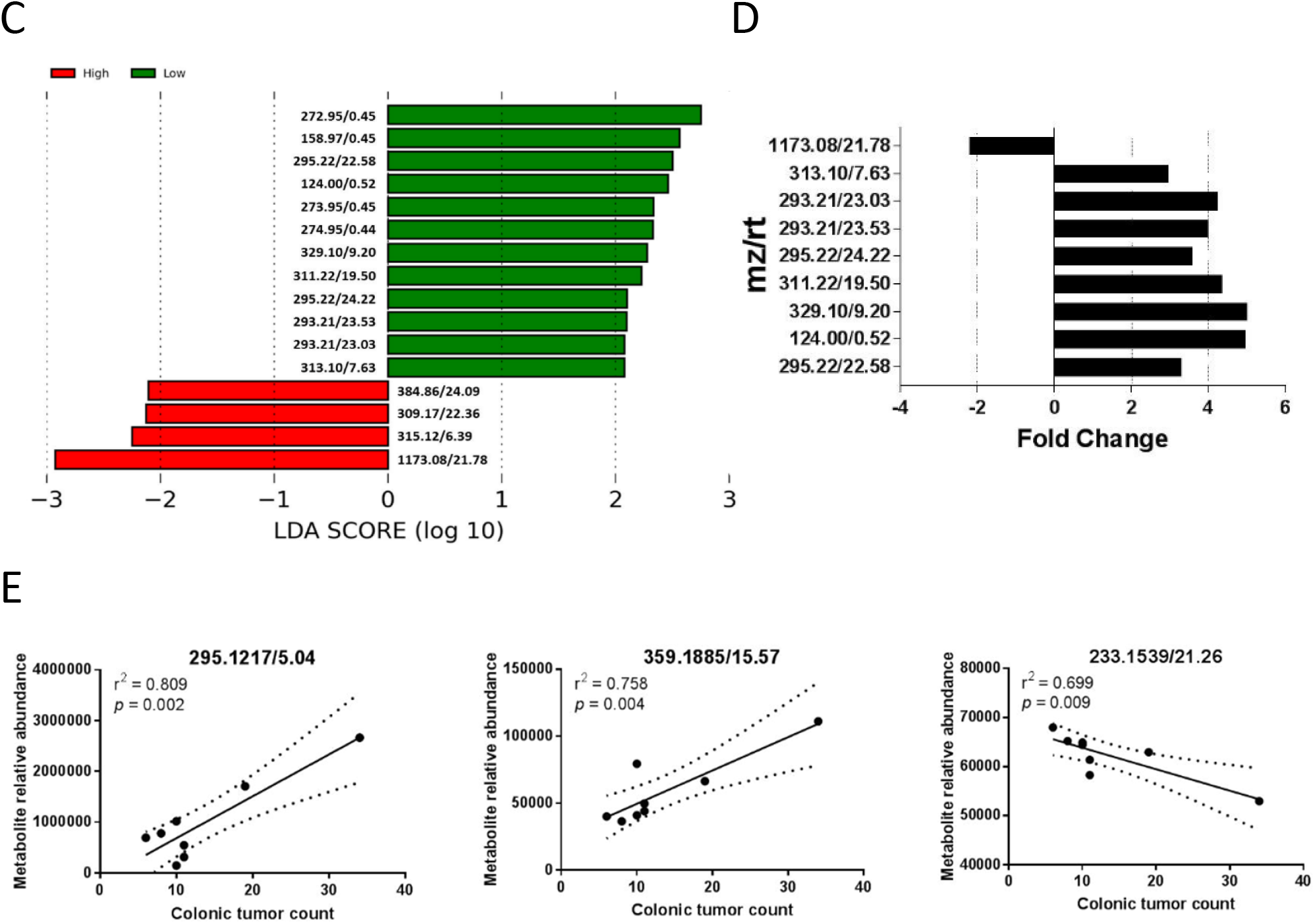
Metabolite features at 1-month of age predict tumor susceptibility and severity. (A) Metabolite features that were significantly different between the high (GM:F344) and low tumor (GM:LEW) groups were used to generate a Heatmap illustrated with the samples along the x-axis and the metabolite features along the y-axis. Hierarchical clustering was performed based on samples and indicates that the GM:F344 samples cluster separately from the GM:LEW group. The fold-change is represented by intensity with red being an increased fold change while blue refers to a decrease. (B) The relative abundance of the top eight differential metabolites between GM:F344 (red dots) and GM:LEW (green squares) are depicted. (C) Linear Discriminant Analysis (LDA) and fold change analysis (D) was used to identify the metabolites driving the Dendrogram tree separation in (A) and differential modulation in high and low tumor groups. (E) Correlation analysis was performed using Pearson’s method to determine positively and negatively correlating metabolites that are associated with increased or decreased tumor multiplicity.

### Bile acid biosynthesis and aspirin-triggered resolvin E biosynthesis pathways are associated with putative fecal metabolomics features

Putative identifications for the differential metabolite features listed are based on the METLIN metabolite library available for public access (Table 1). Based on RMD values, four putative metabolites were classified as steroids while the others were classified as polyphenols, carbohydrates, short chain fatty acids and flavonoids. All putative features identified using UHPLC-MS were subjected to pathway analysis to identify KEGG pathways that were significantly modulated between the two GM profiles. Bile acid biosynthesis (neutral pathway) and aspirin-triggered resolvin E biosynthesis were most differentially affected (Fig.3). The pathway analyses also identified potential genes that may affect or be affected by these putative metabolites (Table 2). The putative identities for the metabolites affecting the bile acid and resolvin E biosynthesis pathways include secondary bile acids such as glycocholate, glycochenodeoxycholate and 7α-hydroxycholest-4-en-3-one (Table 3). We sampled an independent population of Pirc and WT (wildtype) rats at 1-month of age to validate the bile acid and resolvin E biosynthesis pathways as being risk factors for eventual development of adenomas, and to determine if these can be observed in serum. We found that the Pirc animals had elevated serum levels of metabolites related to the bile acid pathway (Supplementary Figure 1A).

**Table 1:** Compound class, RMD and putative identification of metabolites features in the METLIN databases. LC-MS analysis between groups identified several putative metabolites that are listed in the table as mass-charge to retention time ratios. Chemical formulas generated through the Bruker software, along with the calculated relative mass defect and compound classes are also identified. This is additionally supplemented with the putative identification based on the METLIN library.

| Mass-charge/<br>retention time<br>(mz/rt) | Chemical<br>formula | Relative<br>mass<br>defect<br>(RMD) | Compound<br>class | Putative<br>Identification<br>(METLIN ID) |
| --- | --- | --- | --- | --- |
| 124.03995/1.29 | C<sub>6</sub>H<sub>9</sub>O<sub>3</sub> | 322.0737 | Polyphenol | NA |
| 223.0972/6.3 | C<sub>7</sub>H<sub>3</sub>N<sub>2</sub>O<sub>7</sub> | 435.6845 | Carbohydrate | NA |
| 233.15395/21.26 | C<sub>15</sub>H<sub>22</sub>O<sub>2</sub> | 660.2933 | Steroid | 90173 |
| 257.08125/11.52 | C<sub>17</sub>H<sub>8</sub>NO<sub>2</sub> | 316.0479 | Polyphenol | NA |
| 295.1217/5.04 | C<sub>18</sub>H<sub>17</sub>NO<sub>3</sub> | 412.3723 | Carbohydrate | 95663 |
| 297.1127/10.64 | C<sub>18</sub>H<sub>18</sub>O<sub>4</sub> | 379.3173 | Flavonoid | 52682 |
| 359.1885/15.57 | C<sub>24</sub>H<sub>25</sub>NO<sub>2</sub> | 524.7941 | Steroid/SCFA* | 675713 |
| 405.2631/15.12 | C<sub>24</sub>H<sub>36</sub>O<sub>5</sub> | 649.2079 | Steroid | 84737 |
| 527.2822/28.34 | C<sub>13</sub>N<sub>13</sub>O<sub>12</sub> | 535.1973 | Steroid/SCFA* | NA |

\*SCFA – short chain fatty acid; NA – not applicable

**Figure 3.**
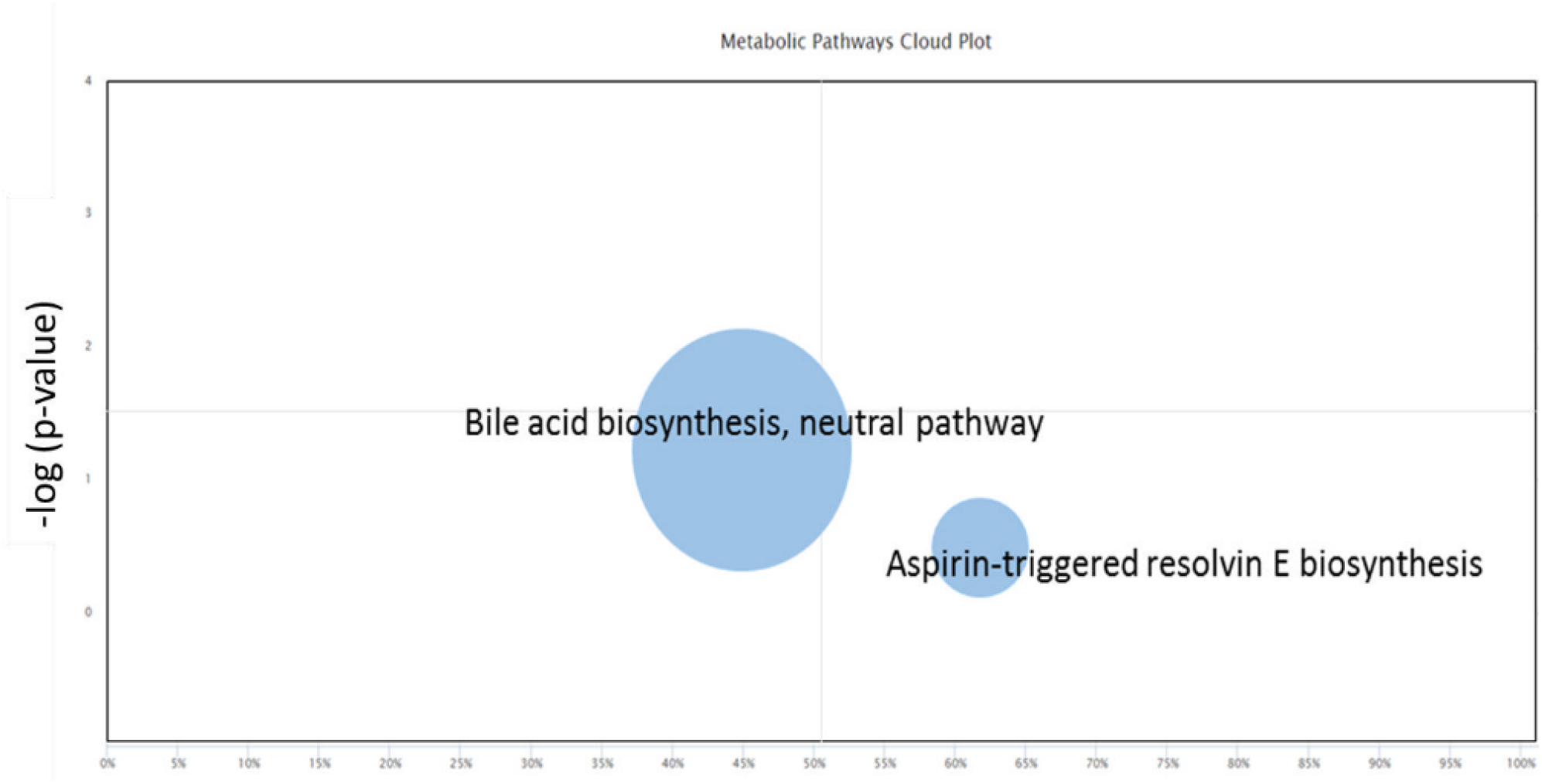
Bile acid biosynthesis and aspirin-triggered resolvin E biosynthesis pathways are most affected by metabolite features. Systems biology analysis, taking into account the differential putative metabolites was performed using the XCMS software. The results showed that bile acid biosynthesis (neutral pathway) and the aspirin-triggered resolvin E biosynthesis were significantly different (*p*< 0.01, Student’s *t*-test) between GM:F344 and GM:LEW. The *p*-value is indicated along the y-axis.

**Table 2:** Genes involved in the bile acid biosynthesis and aspirin-triggered resolving E biosynthesis pathways. The genes listed in the table are part of the putative metabolite pathways differentially regulated between the high and low tumor GM groups. The predicted enzyme activity is listed adjacent to gene names.

| Pathway | Genes | Enzyme activity | FDR-adjusted *p*-value | Group increased in |
| --- | --- | --- | --- | --- |
| Bile acid biosynthesis, neutral pathway | *HSD3B7* | 3 β-hydroxysteroid dehydrogenase type 7 | <u>0.861931</u> | <u>NA</u> |
|  | *ACAA2* | 3-ketoacyl-CoA thiolase, mitochondrial | <u>0.321851</u> | <u>NA</u> |
| | *AKR1D1* | 3-oxo-5-β-steroid 4-dehydrogenase | <u>0.837208</u> | <u>NA</u> |
| | *CYP8B1* | 7-α-hydroxycholest-4-en-3-one 12-α-hydroxylase | <u>0.00627491</u> | <u>GM:LEW</u> |
| | *AMACR* | α-methylacyl-CoA racemase | <u>0.78729</u> | <u>NA</u> |
|  | *BAAT* | bile acid-CoA: amino acid N-acyltransferase | <u>0.0206231</u> | <u>GM:LEW</u> |
|  | *SLC27A5* | bile acyl-CoA synthetase | <u>0.395957</u> | <u>NA</u> |
|  | *SCP2* | chenodeoxycholoyl-CoA synthase | <u>0.85358</u> | <u>NA</u> |
| | *CYP7A1* | cholesterol 7α-monooxygenase | <u>0.0779986</u> | <u>NA</u> |
| | *POR* | cholesterol 7α-monooxygenase | <u>0.093983</u> | <u>NA</u> |
|  | *ACOX2* | peroxisomal acyl-coenzyme A oxidase | <u>0.0756656</u> | <u>NA</u> |
|  | *CYP27A1* | sterol 26-hydroxylase | <u>0.0999591</u> | <u>NA</u> |
|  | *SLC27A2* | very long-chain acyl-CoA synthetase | <u>1</u> | <u>NA</u> |
|  | *CYP2R1* | Vitamin D 25-hydroxylase | <u>0.853166</u> | <u>NA</u> |
| Aspirin-triggered resolving E biosynthesis | *PTGS2* | 18R-hydro(peroxy)-EPE synthase | <u>0.000469693</u> | <u>GM:F344</u> |
|  | *ALOX5* | 5S hydroperoxy HEPE synthase | <u>0.000469693</u> | <u>GM:F344</u> |

**Table 3:**
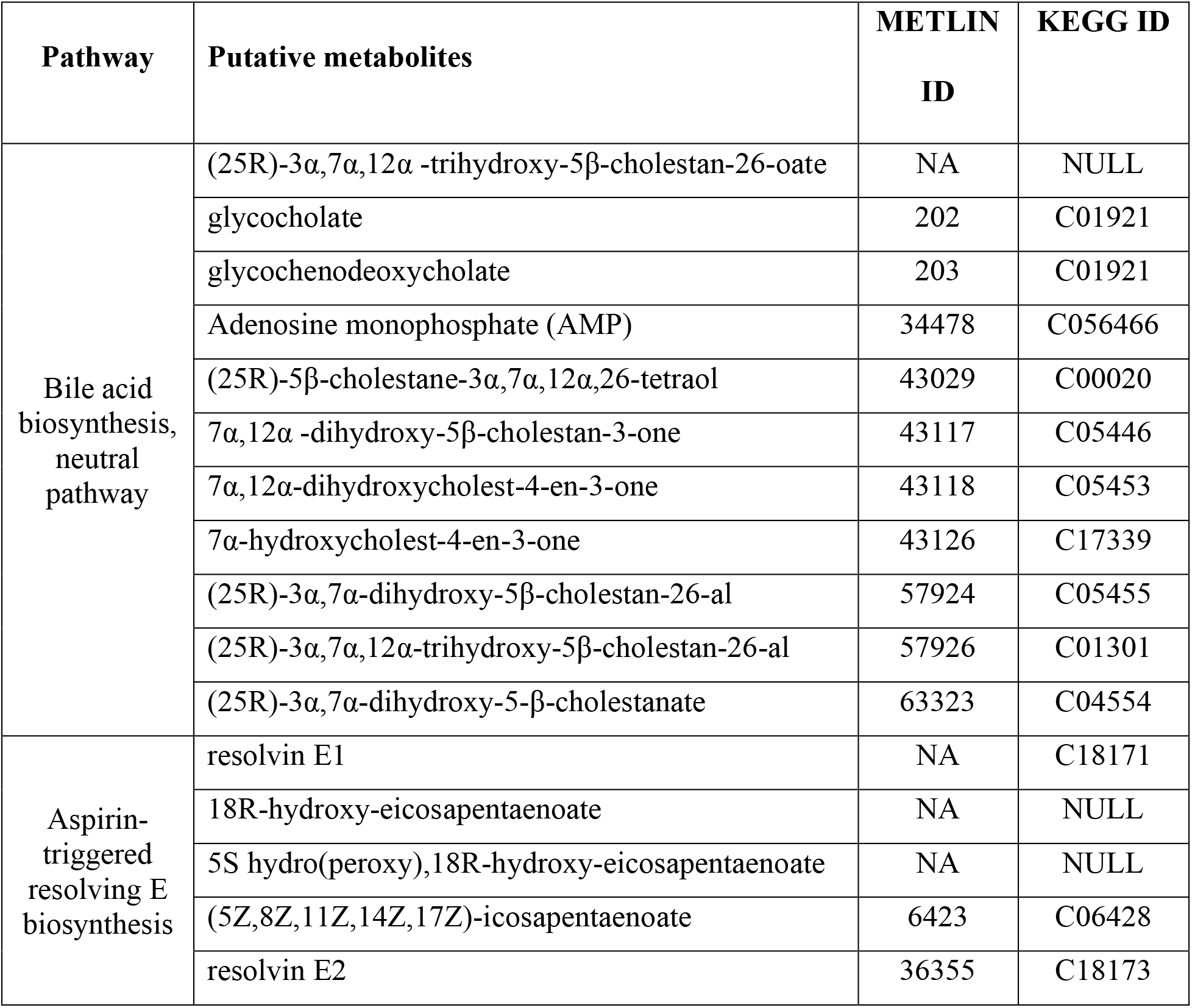
Putative metabolites contributing to bile acid and aspirin-triggered resolving E biosynthesis. The table lists the putative metabolites involved in the bile acid biosynthesis and aspirin-triggered resolvin E pathways. The METLIN and KEGG identification numbers are also listed for testing in the future.

### Gut microbiota is correlated with differences in gene expression in both the normal epithelium and tumor tissues

RNASeq was performed on normal epithelium (NE) and tumor (T) tissues after sacrifice at 6 months of age to determine how the GM may modulate gene expression in isogenic animals. We found that 2173 genes were differentially expressed between GM:F344 and GM:LEW in the normal epithelium tissues (Supplementary Figure 2A and Supplementary Table 2). Additionally, 3406 genes were differentially expressed between adenomas from the two GM profiles (Supplementary Figure 2B and Supplementary Table 2). Clustering analysis (Fig. 4A) showed that the normal epithelium samples separated from the tumor samples, and that each of the tumor and normal epithelial groups separated based on GM profile, i.e. GM:F344 and GM:LEW.

**Figure 4.**
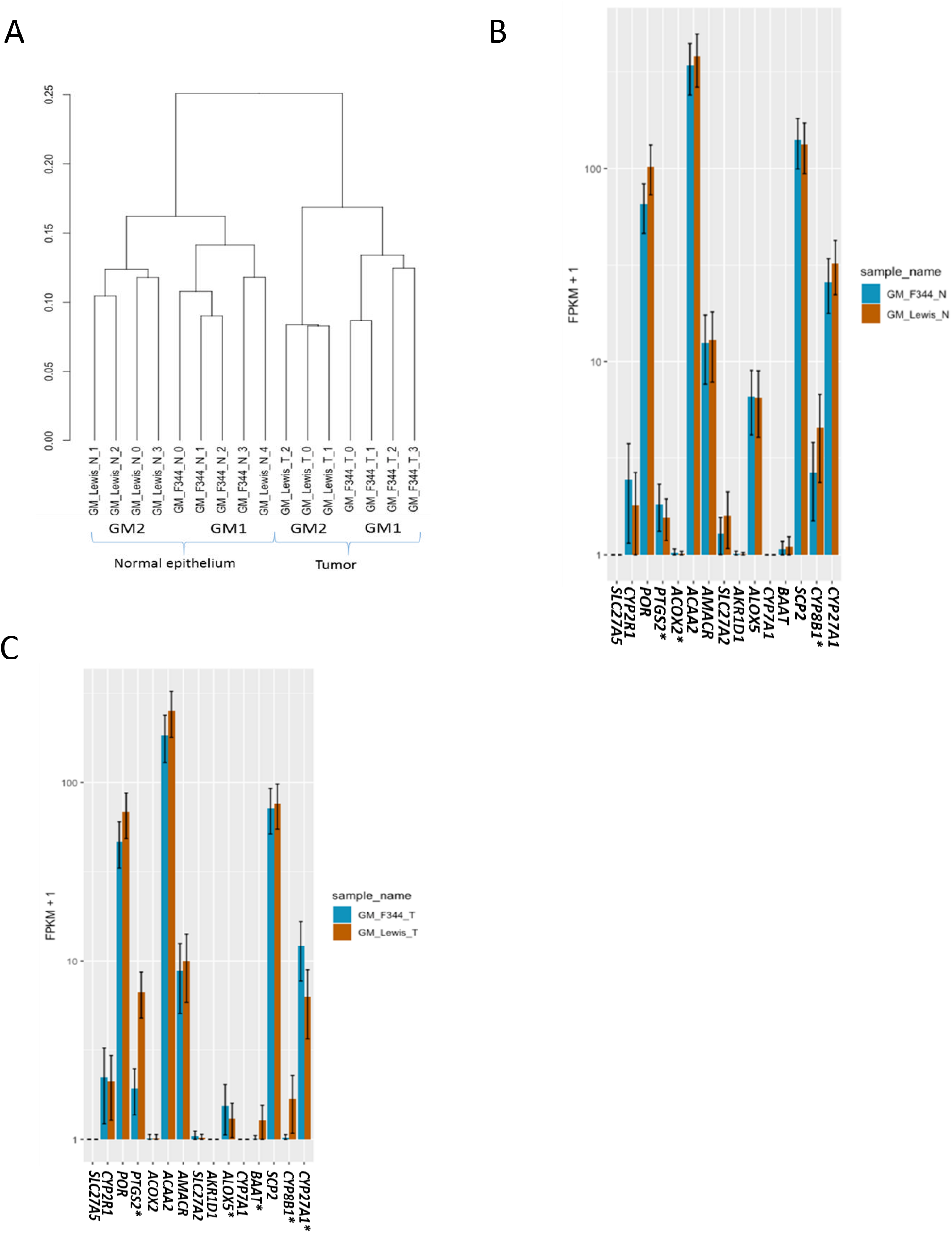
GM correlates with differential gene expression in the normal epithelium and tumor tissues. Ordination and hierarchical clustering (A) analyses were used to determine the relationship of the samples to each other and the groups with respect to the other. Bar plots (GM:F344 – blue, GM:LEW – brown with standard deviation) depicting the relative expression of the genes involved in the pathways affected by the putative metabolites were assessed in the normal epithelium (B) and tumor (C) samples. All the analyses were performed using the cummeRbund package in R. Significantly differential expression with a FDR-adjusted *p*-value less than 0.05 is identified with an asterisk (*) after the gene name.

### Pathway analyses identify potential mechanisms contributing to high and low colonic tumor susceptibility

Pathway analysis using differentially expressed genes found an enrichment in many cancer and metabolism related pathways including the fatty acid and the mucin type-O glycan biosynthesis pathways, with an increased pathway topology in the normal epithelium from the high tumor, GM:F344 group (Supplementary Figure 2C) suggesting a significant role and position of the genes in the pathway. Increased cell cycle, RNA transport, and TCA cycle pathways were also observed in the normal epithelium of GM:F344. Conversely, normal epithelium from the GM:LEW (low tumor) profile showed an increase in apoptotic pathways along with fat digestion and absorption, and calcium signaling pathways (Supplementary Figure 2D).

We determined the expression differences of the genes contributing to the predicted putative metabolic pathways, i.e. bile acid biosynthesis and aspirin-triggered resolvin E biosynthesis (Fig.3, Table 2 and Supplementary Figure 4). We examined the gene expression involved in the resolvin E biosynthesis pathway and found that *PTGS2* was significantly increased in the normal epithelium tissues of the high tumor group (GM:F344) compared to the GM:LEW group (Fig.4B). Interestingly, *PTGS2* was highly elevated in tumor tissues of the low tumor (GM:LEW) group at 6 months of age. We found that *ALOX5* was significantly elevated in the GM:F344 tumor tissues (up 2.5 fold) compared to the GM:LEW group (Fig.4C). Assessing the bile acid biosynthesis genes, we found that *CYP8B1* and *BAAT* were also increased in the tumor tissues of the low tumor (GM:LEW) group compared to Pirc rats in the GM:F344 group (Fig.4C).

We used the differential putative metabolites and differentially expressed genes in the normal epithelium to perform an Integrated Pathway (IP) analysis, while accounting for metabolite, host epithelium expression, and microbiota differences. The synergistic IP analysis suggested that colonic tumor susceptibility is associated with primary bile acid biosynthesis, fatty acid elongation and metabolism pathways. We observed increased pathway topology of unsaturated fatty acid biosynthesis corroborating the role of fatty acids in colonic tumor burden (Fig.5A). To improve the power of our predictive analytical capacity we used canonical correlation analyses to determine the interplay between the OTUs, putative metabolites and the genes identified as differential in the normal epithelium. We found that OTUs such as *Prevotella spp, Desulfovibrio spp, Veillonella parvula* and *Parabacteroides gordonii* are associated with the GM:LEW group in the ordination plot. Similarly, genes *Trim5, Tlr1*, along with *CRABP2, JUNB* and *CNDP2* separate along the axes, based on their relationship with either GM:F344 or GM:LEW. While a putative metabolite identified as vigabatrin correlated with GM:LEW, the other metabolites detected clustered with GM:F344 in the analysis (Fig.5B).

**Figure 5.**
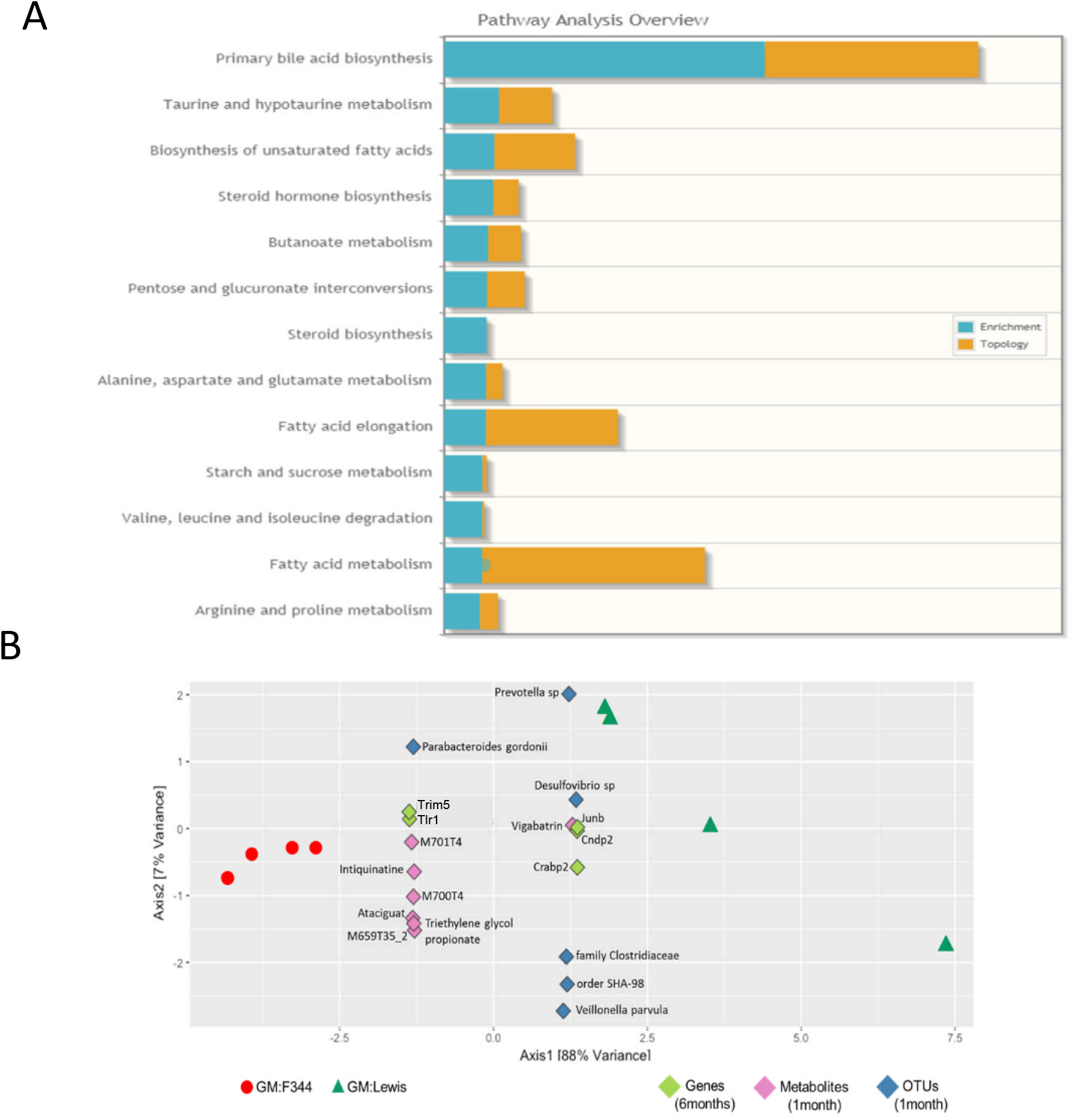
Pathway and correlation analyses identify potential mechanisms, differential factors contributing to low, and high tumor susceptibility. (A) Integrated pathway analyses depicts pathways enriched and their topology, contributing to the variability in tumor phenotype observed as an effect of the genes and metabolites. (B) Sparse canonical correlation analysis incorporating the genes, metabolites and OTUs contributing to disease susceptibility in GM:F344 (red dots) and GM:LEW (dark green triangles) were analyzed in R, using the structSSI CRAN package. Metabolites, genes and OTUs are shown as diamonds in purple, light green and blue. Axis-1 demonstrated an 88% separation between GM:F344 and GM:LEW.

## Discussion

Colon cancer etiology has been addressed for decades from the perspective of host gene expression and its effect on disease susceptibility. Studies have also addressed the metabolome associated with tumorigenesis separately or in conjunction with the microbiome or the transcriptome. However, the majority of these studies are retrospective, i.e. after disease onset in patients, raising the question of whether the microbiome, metabolome and transcriptome are merely responding to disease, or causative of tumor development. We present for the first time the integration of three ‘omics’ strategies to understand tumor susceptibility in a preclinical rat model of human colon cancer. Multi-omics investigations included integrated pathway analysis combining the metabolomics and transcriptomics data, identifying potential biomarkers for disease risk from fecal samples as early as 1 month of age.

Previously, we reported that differential commensal GM altered the susceptibility of isogenic Pirc rats rederived onto surrogate dams with distinct gut microbiota (7). We now report that the altered GM profile correlates with differential metabolite features representative of the high tumor, GM:F344 and the low tumor, GM:LEW profiles. Some of the differential putative metabolites have identities in the METLIN database, facilitating future testing of these compounds and their influence on tumorigenesis. We remarkably found that Pirc rats with fewer adenomas (<9, average) differentially clustered from animals with more than 19 adenomas, irrespective of the GM. These metabolite data were prognostic at 1-month of age, substantially prior to the onset of visible adenomas and physiological signs of disease in Pirc rats, suggesting a role for the metabolites in the initiation of early adenomas. Due to the inadequacy of compound libraries in their current state, we could not establish accurate identities of the compounds using tandem MS spectra. However, further investigation including advanced methods such as UHPLC-MS-SPE-NMR (solid phase extraction and nuclear magnetic resonance) could elucidate the identity of these metabolites (17, 18). This information will be used going forward as training datasets for neural network or machine learning algorithms with the objective of establishing a pre-tumorigenesis dataset to identify at-risk populations based on metabolite features (19, 20).

Similar to the metabolomic analysis, we demonstrated here for the first time that congenic Pirc rats show differential gene expression depending on the GM they harbor. We found that *PTGS2* was significantly elevated in the normal epithelium in the GM:F344 group suggesting that the gut microbiota likely has a role in the differential expression of this gene. *PTGS2* is an integral gene in the cyclooxygenase-2 (*COX2*) mechanism and has been associated with increased colonic tumor burden (21). We also found increased *ALOX5* expression in GM:F344 tumors, which is associated with increased proliferation and invasion of colonic tumors (22). Interestingly, the *COX2* mechanism highlighted in our study is based on metabolic pathways obtained from fecal samples collected at 1 month of age suggesting that the microbiome may be mediating low level long-term effects through the metabolites.

Increased bile acid exposure in the gastrointestinal tract is a known factor for GI cancers and were proposed as carcinogens in as early as 1939 (23, 24), with secondary bile acids reported to be significantly increased in serum from patients with colonic adenomas (25). In accordance to these reports, we found that the bile acid biosynthesis was elevated in the high tumor (GM:F344) group. Using an Integrated pathway analysis based on differential metabolites and the differentially expressed genes in the normal epithelium, we found primary bile acid biosynthesis as one of the principal contributors to the variability in disease susceptibility observed in GM:F344 or GM:LEW rats.

We concurrently used sparse canonical correlation analysis to integrate the microbiome, metabolome and the transcriptome to identify potential features associated with disease phenotype and susceptibility. This approach was crucial to increase our confidence of prognostic feature detection as the metabolite identifications have not yet been verified through more-advanced methods such as NMR. Based on this approach we found that several putative metabolites (M701T4, M700T4, Triethylene glycol propionate) and two genes (*Trim5* and *Tlr1*) correlated with the increased tumor phenotype of the GM:F344 group. Interestingly, TRIM5 (tripartite motif 5) and TLR1 (toll-like receptor 1) are both significant contributors to the innate immune system. TRIM proteins in humans are E3 ubiquitin ligases are known to contribute to inflammatory cytokine production (26). Like the TRIM5 proteins, toll-like receptors (TLRs), activate the adaptive immune response in cancers. Lu *et al* recently showed that TLR1 expression was elevated along with other TLRs in colorectal cancer (CRC) patients and cancer cell lines. The elevated expression was associated with the simultaneous increase in expression of pro-inflammatory genes such as IL-6 (interleukin-6) which has been identified as a potential marker for diagnosis and predicting severity of CRC (27–29). The association of these genes with the high tumor (GM:F344) group suggests that the adenoma development in our model is mediated via inflammation and/or that their expression is a response to the resident taxa. However, due to the putative identities and nomenclature of the metabolites future studies will be required to validate their association with increased disease susceptibility. We also found that the relative abundance of OTUs (*Prevotella, Desulfovibrio*, and *Parabacteroides spp*) (30–32), previously linked with reduced colon cancer (7) were simultaneously associated with the low tumor susceptibility (GM:LEW) group. Using bacterial supplementation methods, the importance of these taxa and their role in colon cancer will also need to be carefully addressed going forward.

We found that the microbiome, metabolome and transcriptome play a large role in the etiology of colon cancer, with the GM influencing the other two components enormously. Assimilating these omics strategies has led to the discovery of several targets in all three systems that in the future could be used for screening, and potentially therapeutic interventions. Our data and approach could enhance precision medicine both in a diagnostic and prognostic manner in the future. More importantly, we demonstrated that the complex GM is an important factor that needs to be defined or controlled for in all studies examining drug or therapeutic interventions because of the altered metabolic profile and the host response.

## Methods

### Animal husbandry and housing

Pirc rats were generated by crossing male, F344/Ntac-*Apc^+/am1137^* Pirc (RRRC:782) rats with wild-type female rats obtained commercially from Envigo Laboratories (Indianapolis, IN), i.e. F344/NHsd. All animals were group housed, prior to time of breeding on ventilated racks (Thoren, Hazleton, PA) in micro-isolator cages. Cages were furnished with corn cob bedding and were fed irradiated 5058 PicoLab Mouse Diet 20 (LabDiet, St. Louis, MO). Rats had *ad libitum* access to water purified by sulfuric acid (pH 2.5-2.8) treatment followed by autoclaving. Prior to breeding fecal samples were collected from both the breeders using aseptic methods and banked at −80 °C.

All procedures were performed according to the guidelines regulated by the Guide for the Use and Care of Laboratory Animals, the Public Health Service Policy on Humane Care and Use of Laboratory Animals, the Guidelines for the Welfare of Animals in Experimental Neoplasia, and were approved by the University of Missouri Institutional Animal Care and Use Committee.

### Experimental design

We used previously collected fecal and tissue (normal epithelium or tumor) samples from F344-*Apc^+/am1137^* PIRC rats generated through complex microbiota targeted rederivation (CMTR) as described by Ericsson *et al* (33). These previously banked samples were used in this study to assess how the GM affects the metabolome and transcriptome (Fig.1). Briefly, fecal samples collected from animals aseptically at 1 month of age and prior to onset of observable colonic tumor phenotype for metabolomics were collected into and immediately snap frozen with liquid nitrogen and stored at −80 °C until processing for metabolomics. At 6 months of age, animals were sacrificed post-disease onset, confirmed through colonoscopies described previously (21). Tumor (T) and adjacent normal epithelium (NE) tissues were collected into cryovials aseptically, flash-frozen and stored at −80 °C.

### Genotyping and animal identification

Pups were ear-punched prior to weaning at 18 days of age using sterile technique. DNA was extracted using the “HotSHOT” genomic DNA preparation method previously outlined (34). Briefly, ear punches were collected into an alkaline lysis reagent (25 mM NaOH and 0.2 mM EDTA at a pH of 12). The ear clips were heated at 90 °C on a heat block for 30 minutes, followed by addition of the neutralization buffer (40 mM Tris-HCl) and vortexing for 5 seconds. Obtained DNA was used for a high resolution melt (HRM) analysis as described previously (7).

### Serum collection

Pirc and WT rats were anaesthetized with isoflurane at 1-month of age. 0.5 ml of blood was drawn aseptically via the jugular vein and the serum was collected by precipitating the cells at 10,000 g for 10 minutes. The collected serum was centrifuged again at 16,000 for 5 minutes to remove any lysed debris or cells, and then stored at −80 °C until further processing in glass vials.

### Ultra-high performance liquid chromatography and mass spectrometry (UHPLC-MS)

Fecal samples were lyophilized at −20 °C using 0.1 millibar of vacuum pressure, following which dried samples (30 mg) were extracted sequentially for both UHPLC-MS and GC-MS. The dried samples were first treated with 1.0 mL of 80% MeOH containing 18 μg/mL umbelliferone, sonicated for 5 minutes and centrifuged for 40 minutes at 3000 g at 10 °C. 0.5 mL of supernatant was used for UHPLC-MS analysis after a subsequent spin at 5000 g at 10 °C for 20 minutes and transferring 250 μL of the sample into glass autosampler vials with inserts. For GC-MS (Gas Chromatography-Mass Spectrometry) analyses of primary polar metabolites, 0.5 mL water was added the remaining extract used above for the UHPLC preparation, sonicated for 5 min, extracted for 30min, and centrifuged at 3000 g. 0.5 mL of the polar extract was subsequently dried under nitrogen and derivatized using previously established protocols (35). Briefly, N-Methyl-N-(trimethylsilyl) trifluoroacetamide (MSTFA) with 1 % TMCS (2,2,2-Trifluoro-N-methyl-N-(trimethylsilyl)-acetamide, Chlorotrimethylsilane) was used to derivatize the polar metabolites, after treatment with methoxyamine-HCl-pyridine. UHPLC-MS analyses were performed on a Bruker maXis Impact quadrupole-time-of-flight mass spectrometer coupled to a Waters ACQUITY UPLC system. Separation was achieved on a Waters C18 column (2.1x 150 mm, BEH C18 column with 1.7-μm particles) using a linear gradient composed of mobile phase A (0.1% formic acid) and B (B: acetonitrile). Gradient conditions: B increased from 5% to 70% over 30 min, then to 95% over 3 min, held at 95% for 3 min, then returned to 5% for re-equilibrium. The flow rate was 0.56 mL/min and the column temperature was 60 °C.

Mass spectrometry was performed in the negative electrospray ionization mode with the nebulization gas pressure at 43.5 psi, dry gas of 12 l/min, dry temperature of 250 °C and a capillary voltage of 4000V. Mass spectral data were collected from 100 and 1500 m/z and were auto-calibrated using sodium formate after data acquisition.

Metabolites that were significantly different between each group and that contributed to the Dendrogram separating low and high tumor animals were selected for targeted tandem MS (MS/MS) analysis. MS/MS spectral data were collected using the following parameters: MS full scan: 100 to 1500 m/z; 10 counts; active exclusion: 3 spectra, released after 0.15 min; collision energy: dependent on mass, 35 eV at 500 Da, 50 eV at 1000 Da and 70 eV at 2000 Da. Mass spectra were calibrated using sodium formate that was included as a calibration segment towards the end of the gradient separation.

### Metabolomics Data Processing

For UHPLC-MS data, the mass spectral data were first calibrated using sodium formate and converted into net.CDF file format for processing using XCMS (ref: https://www.ncbi.nlm.nih.gov/pubmed/16448051) that included peak detection, deconvolution, alignment and integration. The signal intensities were then normalized to that of the internal standard umbelliferone (abundance of metabolite/abundance of umbelliferone × 100%) and used for statistical analysis. MS/MS spectra were searched against our custom spectral library (36) and the Bruker libraries (https://www.bruker.com/products/mass-spectrometry-and-separations/metabobase-plant-libraries/), MassBank of North America (MoNA, http://mona.fiehnlab.ucdavis.edu/), mzCloud (https://www.mzcloud.org/) for confident or putative identifications. Multivariate statistical analysis such as principal component analyses (PCA) and ANOVA was performed using MetaboAnalyst (http://www.metaboanalyst.ca/) after pre-treatments of the data, i.e. normalization to sum, log transformation, and auto scaling.

### Fecal DNA extraction, 16S library preparation and sequencing

Fecal samples were pared down to 70 mg using a sterile blade and then extracted using methods described previously (37). Amplification of the V4 hypervariable region of the 16s rDNA and sequencing was performed at the University of Missouri DNA core facility (Columbia, MO) as previously described (37).

### Normal epithelium and tumor tissue collection

All animals were humanely euthanized with CO_2_ (carbon di-oxide), administration and necropsied at sacrifice. The small intestine and colon from the rats were placed on to bibulous paper and then splayed opened longitudinally by cutting through the section. Using a sterile (Feather, Tokyo, Japan) scalpel blade normal colonic epithelium tissues were scraped from the top, middle and distal regions of the colon. Tumors in the same locations were collected by resecting half of the total tissue. All tissues were flash-frozen in liquid nitrogen and stored at −80 °C. Remaining intestinal tissues were then fixed overnight in 10% formalin, which was then replaced with 70% ethanol for long term storage until adenoma counting was performed.

### Tumor counts and measurements

Tumor counts were estimated as previously described using a M165FC (Leica, Buffalo Grove, IL) microscope at 0.73X magnification (7). Briefly, the small intestine and colonic tissues were laid flat in a large petri dish (Sycamore Life Sciences, Houston, TX) and covered with 70% ethanol (ThermoFisher Scientific, Waltham, MA) to prevent tissue drying. Biologic forceps (Roboz Surgical Instruments Co., Inc., Gaithersburg, MD) were used to gently count polyps observable under the objective. Tissues were kept hydrated throughout the entire process. Tumor sizes were measured using the Leica Application Suite 4.2, after capturing post-fixed images as previously described (7).

### RNASeq and bioinformatics analysis

Normal epithelium and tumor tissue samples were collected upon necropsy at 180 days of age and were extracted using the Qiagen AllPrep DNA/RNA mini kit (Qiagen, Germantown, MD) after pre-processing using the QIAshredder (Qiagen, Germantown, MD) columns to extract total RNA. The quality of RNA was then assessed using the Experion RNA StdSens analysis kit (Bio-Rad, Hercules, CA). Based on the RNA-quality index (RQI), 18S and 28S peaks in the chromatogram, samples were classified into high (>9), medium (7> or <9) or poor quality (>6). Except for one sample (normal epithelium from rat 044, i.e. 044_N), all other samples were of medium or higher RQI. Total RNA was used for poly-A selection and Illumina TruSeq paired-end library preparation following manufacturer’s protocols. 75 bp (basepair) paired-end reads were sequenced on the Illumina MiSeq (38) platform to an average of depth of 50 x 10^6^ reads per sample. All samples were processed at the same time and sequenced on a single lane, to avoid batch effects.

Sequence read alignment was done using Tophat from the Tuxedo protocol as outlined in the original publication (39). To remove adaptors and low-quality reads, Trimmomatic v.0.32 was used with standard settings (40), and then aligned to the Rat genome (Rnor_6.0) (download from: ftp://igenome:G3nom3s4u@ussd-ftp.illumina.com/Rattus_norvegicus/NCBI/RGSC_v3.4/Rattus_norvegicus_NCBI_RGSC_v3.4.tar.gz on May 24th, 2017) using Tophat2 v2.0.12 with default settings. The aligned reads were sorted with SAMtools v1.3, followed by HTseq v0.9.1. Differential gene expression was then estimated using the DESeq2 v1.18.1 in R v3.4 (41). Read count distributions in the normal epithelium and tumor tissues were found to be bimodal, with genes being identified as significant based on an FDR-adjusted *p*-value of < 0.05 and with a fold-change of at least 1.5-fold. Pathway analyses were performed on the top 100 significantly up-regulated genes in either GMs, i.e. GM:F344 or GM:LEW. Pathway over-representation analyses were based on hypergenometric distribution to determine the statistical significance of a particular gene to an over-represented pathway. Topology analysis was also performed using the degree centrality method and the gene-centric Integrated Pathways (IP) module of Metaboanalyst v3.0 (42). For the IP analysis, the list of differentially abundant putative metabolites and differentially expressed genes between the high and low tumor normal epithelium samples were used as input. Briefly, the genes and metabolites are mapped on to KEGG pathways to identify pathways that are over-represented and show increased topology, the latter signifying the relative importance to a given pathway. Enriched pathways based on this analysis were selected using a FDR-adjusted *p*-value of < 0.05. A similar analysis was performed for both the NE and T samples. We also used the sparse Canonical Correlation Analysis (sCCA) (43) to identify the genes, OTUs and putative metabolites that contribute to the covariation observed in disease susceptibility. This analyses allowed for detection of entities correlating with the high and low tumor phenotypes. Briefly, we *log*-transformed the bacterial abundance, retaining only the OTUs assigned to the Genus-level due to inflated zero abundance levels in the abundance tables. No transformation was employed for the putative metabolite abundance and gene count tables. Thereafter, the sCCA was performed using the procedure described by Callahan *et al*. (44).

### Metabolomics analyses

Mass spectral data from each sample were converted into netCDF formatted files and processed with XCMS to generate lists of mass features and their intensities (45). An average of 499 peaks were found per sample. Peaks appearing in less than a quarter of the samples in each group were ignored. 175 variables were removed for threshold 25 percent, i.e. appearance of peaks in greater than 25% of the samples per group. Variables with missing values were replaced with a small value (0.0000001) for statistical analysis purposes. The data were then normalized to sum, transformed using Log normalization and auto-scaled to ensure maximum-possible binomial distribution. The number of samples, raw peak numbers observed and the final peak list used for each sample processed are described in Supplementary Table 1.

Statistical analyses were performed based on a threshold of 2, for the fold-change analysis, with values displayed in the log-scale to observe both the up-regulated and downregulated features in a symmetrical way. Principal Component Analysis (PCA) was performed using the *prcomp* package in R using the *chemometrics.R* script (46). NMDS (non-metric dimensional scaling) is another method for ordination and was performed using the *vegan* package in R (47).

Hierarchical clustering analysis was performed using the Euclidean distance measure using the *Ward* algorithm (to minimize the sum of squares of any two clusters, potentially separating only if large differences exist between groups) and displayed as a Dendrogram using the *hclust* function in the *stat* package in R. To determine the metabolites contributing to the separation and rooting of the hierarchical clusters, the samples irrespective of GM were re-classified into those with ‘high’ or ‘low’ tumors and a linear discriminant analysis (LDA) was performed using the LEfSe module on a high-computing Linux platform (48) with a LDA score of log_10_2 or greater being significantly differential metabolites between the high and low tumor groups.

### Statistical analyses and figures

All other statistical analyses were performed using Sigmaplot 13.0 (Systat Software, San Jose, CA) and graphing for figures (except Fig.1) was prepared through GraphPad Prism version 7 for Windows (GraphPad Software, La Jolla, CA). *p*-values were set to identify significance at a value less than 0.05, unless otherwise described or indicated. Correlations were performed using the linear regression module available through GraphPad Prism v7.

### Ethics Statement

The study reported here was conducted in accordance with the guidelines established by the Guide for the Use and Care of Laboratory Animals and the Publish Health Service Policy on Human Care and Use of Laboratory Animals. All studies and protocols (#6732 and #8732) were approved by the University of Missouri Institutional Animal Care and Use Committee.

### Availability of Data and Material

The RNAseq raw and processed data files generated and analyzed during the current study are available at the NCBI Gene Expression Omnibus (GEO) database repository under the accession ID# GSE120934. The associated BioProject and SRA (sequence read archive) numbers are PRJNA495060 and SRP164571 respectively. The raw data for the metabolomics analyses is hosted through the Metabolomics Workbench on the NIH Metaboloics Data Repository under the DataTrack ID #1539 for public access.

## Supporting information

Supplemental RNASeq

## Author contributions

Experiments were conceived and designed by Susheel Bhanu Busi, Lloyd W. Sumner and James Amos-Landgraf. Zhentian Lei and SB performed the experiments with reagent and instrumental contributions from ZL and LWS. Data was analyzed by SB and ZL. All authors contributed to writing the paper.

## Acknowledgements

The authors wish to acknowledge Miriam Hankins, Marina McCoy, Rebecca Schehr, Aaron Ericsson and Elizabeth Bryda for assistance with fecal collection; Nathan Bivens and the MU DNA Core for assistance with 16S rDNA and RNAseq experiments; Bill Spollen and the MU Informatics Research Core Facility for assistance with software installation for data analysis; Rat Resource and Research Center; MU Office of Animal Resources and their staff for assistance with animal husbandry.

## Funding

This research was funded by grants from the University of Missouri to Dr. James Amos-Landgraf (Startup-funding; 2012 - present) and the MU Research Board grant awarded to Dr. Amos-Landgraf (2017). The MU Metabolomics Center is supported by the University of Missouri Office of Research, and the Sumner lab is supported by NSF Awards 1340058, 1139489, 1126719. The Sumner lab is also supported by Bruker Daltonics, Gmbh.

## Conflict of interest statement

The authors declare that the research was conducted in the absence of any commercial or financial relationship that could be construed as a potential conflict of interest.

## (Figures including Supplementary) 7.24.20

**Supplementary Figure 1.**
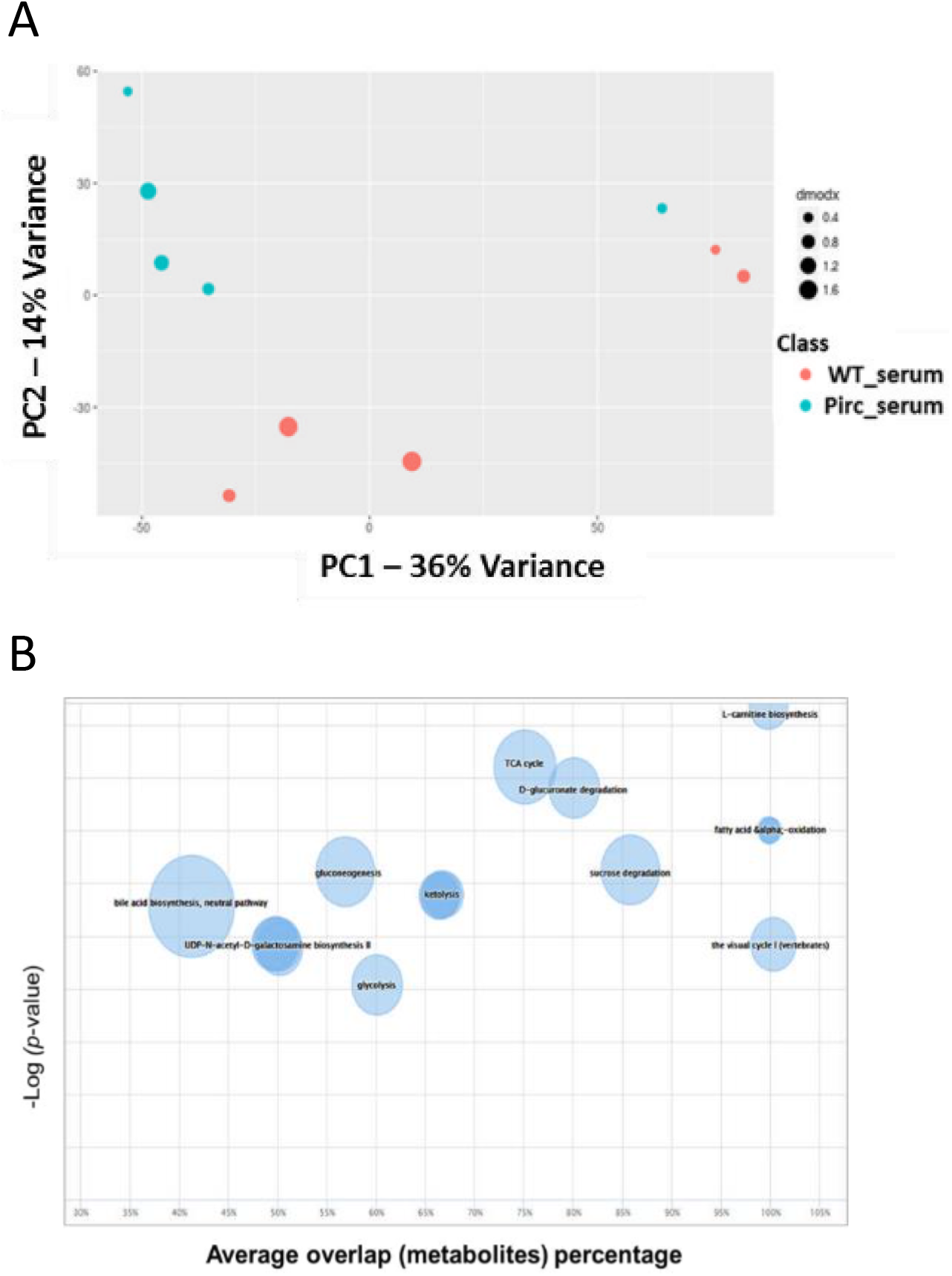
Serum metabolomics profiles and pathway analyses in Pirc and WT rats. Serum metabolomics profiles and pathway analyses in Pirc and WT rats Serum samples collected from Pirc and WT rats at 1-month of age used for LC-MS analysis indicated differential metabolomics profiles (A) including the regulation of bile acid biosynthesis, L-carnitine biosynthesis and fatty acid alpha-oxidation as potential pathways (B), contributing to phenotypic differences.

**Supplementary Figure 2.**
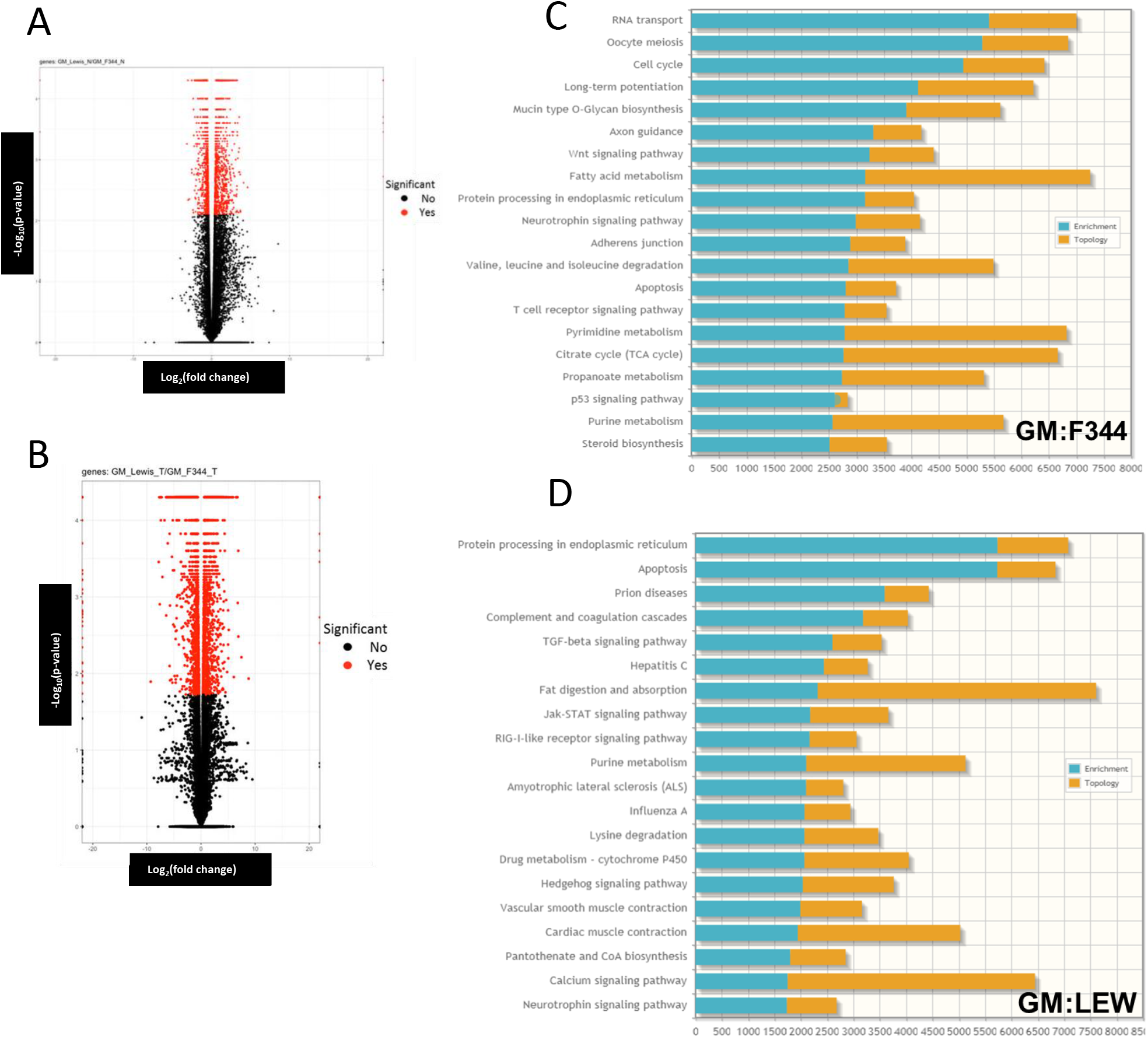
Differentially expressed genes (DEGs) altered due to GM in the normal epithelium. Volcano plot analysis was performed on the differential gene expression in both the normal epithelium (A) and the tumor (B) samples from GM:F344 and GM:LEW. Fold change and *p*-values are established along the x- and y-axes. All genes with a fold change of at least 2, and FDR-corrected *p*-value were used for further analysis. Pathway analyses based on the expression of upregulated genes in the normal tissues of the GM:F344 (C) and GM:LEW (D) groups was used to identify potential pathways and mechanisms contributing to the low and high tumor susceptibility. Enriched pathways are indicated in blue, while the topology, i.e. the importance of the pathway to the overall phenotype observed is shown in yellow. Integrated pathways (IP) analysis incorporating the differentially expressed genes and the putative metabolites, significantly different between GM:F344 and GM:LEW, was performed.

**Supplementary Figure 3.**
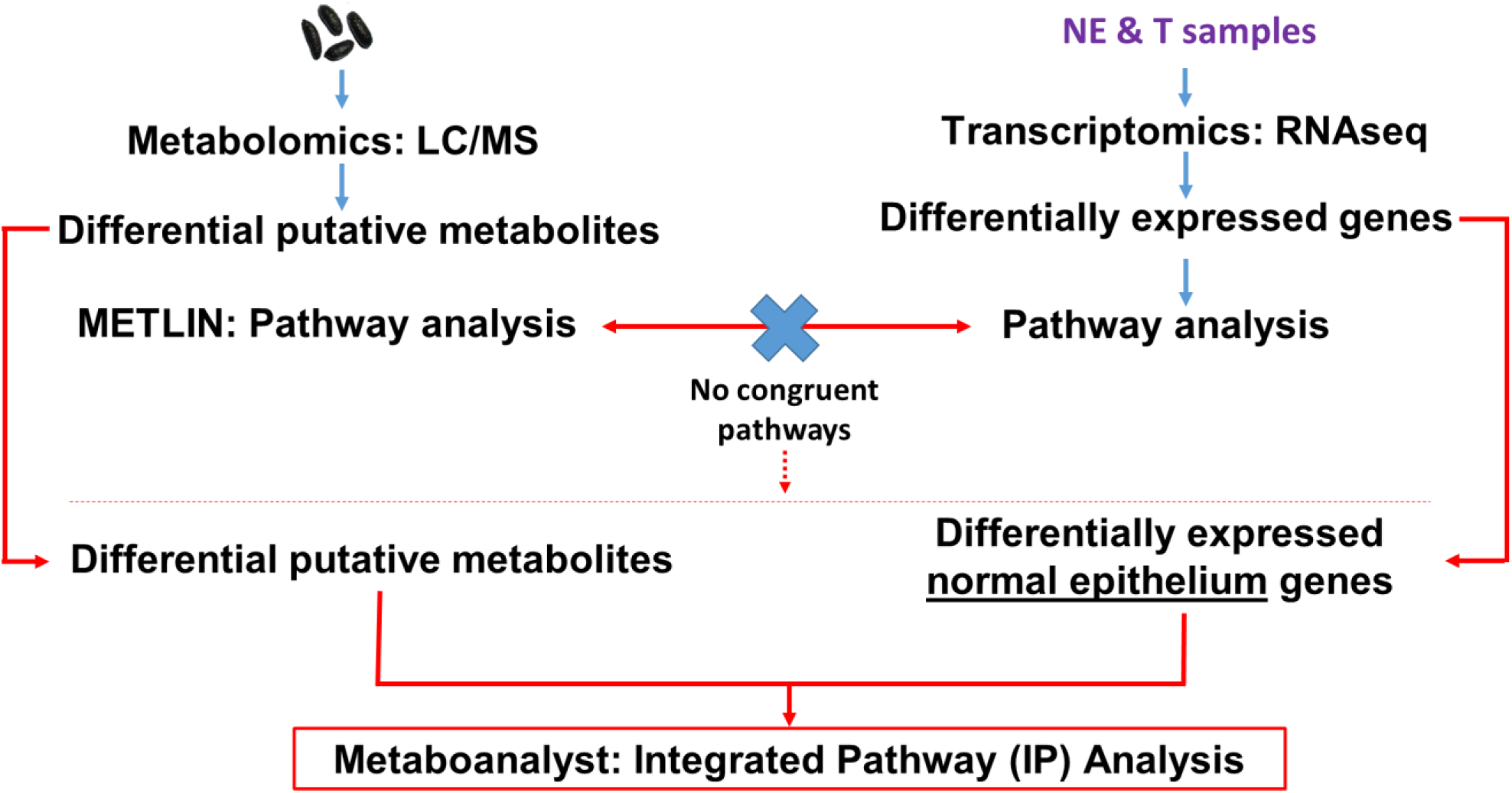
Analysis flowchart. Flowchart describing the processing and analysis methods employed for the fecal and tissue samples. Fecal samples were processed for metabolomics. Normal epithelium (NE) and tumor (T tissues were used for transcriptomics (RNASeq) analysis. Integrated Pathway analysis was subsequently performed using the differential putative metabolites and the differentially expressed genes from the normal epithelium of the high and low tumor groups.

**Supplementary Figure 4.**
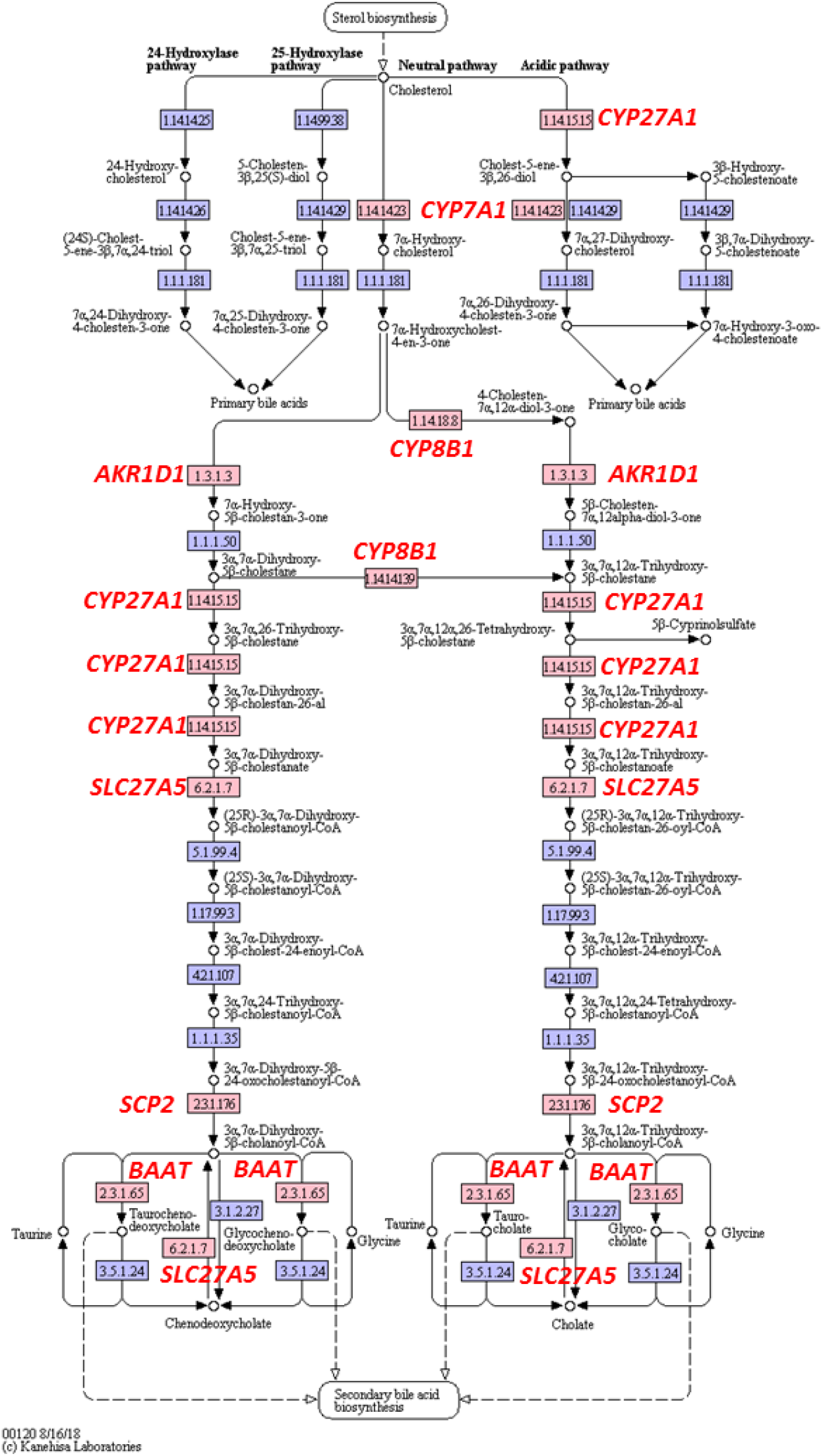
Bile acid biosynthesis pathway. Genes identified via metabolomics and RNAseq analysis, contributing to the bile acid pathway analyses are identified by highlighting corresponding locations in the KEGG pathway. The pathway was built using the KEGG pathway mapper tool from www.genome.jp/KEGG

## Notes

https://www.ncbi.nlm.nih.gov/geo/query/acc.cgi?acc=GSE120934

https://www.metabolomicsworkbench.org/data/DRCCMetadata.php?Mode=Study&StudyID=ST001075

### Competing Interest Statement

The authors have declared no competing interest.

