## Supplemental Tables for "Integrated metabolome and transcriptome analyses provide insight into colon cancer development by the gut microbiota"

**Supplementary Table 1: Summary of data processing results**

| Samples | Peaks (raw) | Missing/Zero | Peaks (processed) |
|---|---|---|---|
| SB1 | 536 | 126 | 246 |
| SB2 | 423 | 167 | 246 |
| SB3 | 528 | 148 | 246 |
| SB4 | 498 | 149 | 246 |
| SB5 | 500 | 129 | 246 |
| SB6 | 537 | 100 | 246 |
| SB7 | 491 | 129 | 246 |
| SB8 | 478 | 169 | 246 |
| SB9 | 486 | 151 | 246 |
| SB10 | 512 | 127 | 246 |
| SB11 | 493 | 159 | 246 |
| SB12 | 501 | 148 | 246 |
| SB13 | 479 | 155 | 246 |

The raw peaks obtained via XCMS for each individual samples analyzed through LC-MS is shown with an average peak abundance in the samples being 497. The number of missing or zero peaks for each sample along with the number of peaks processed for analysis based on the cutoff established in the Methods sections are listed. The raw data for the metabolomics analyses is hosted through the Metabolomics Workbench on the NIH Metaboloics Data Repository under the DataTrack ID #1539 for public access.

**Supplementary Table 2: Differentially expressed genes in the normal epithelium and tumor tissues of GM:F344 and GM:LEW**

Busi_RNASeq_Differential_Genes.xlsx
